# Interpenetration between a polymer brush and a polymer star at thermal equilibrium: A theoretical approach

**DOI:** 10.1101/489559

**Authors:** Mike Edwards

## Abstract

By means of the density functional theory framework I tackle the long-standing problem of a polymer star interpenetrating with a polymer brush at thermal equilibrium. Remarkably, the star is repelled to the outside of the brush once it sucks into the brush. It turns out that there could be a highly fluctuating region at the brush edge. The highly fluctuating region would be responsible for discontinuous absoption transitions by brushes. However, up to an small interpenetration length, below which asphericity of the star is maintained, the star gets collapsed by sucking more and more into the brush.

## INTRODUCTION

Polymers are large chain-like macromolecular structures that have been made from repeated atomic or molecular sub-units called monomers^1,5^. Typically, number of monomers per chain i.e. the degree of polymerization is between 10^4^ and 10^5^. The connectivity among monomers is made through covalent bond in which two atoms share their valence electron. The covalent bond is strong enough to prevent chains wrap apart. Polymers can be found in nature with a wide variety of architectures. In this article, I concentrate on two particular polymeric structures with respect to their importance in biology, nanotechnology, medicine and so on. Here, I am going to address, theoretically, polymer brushes and polymer stars as well as their equilibrium properties when they interpenetrate into each other. Polymer brushes are formed when linear chains are grafted to (from) a substrate such that the steric repulsion between nearby chains stretches all chains in perpendicular direction^2^. Brush-like structures are found in every biological systems where there is a fluffy surface covered by polymers. For instance, glycol on cell membranes or aggregan in synovial joints optimize blood flow or lubrication, respectively. Interested reader may see Ref.^4^ for a recent review on theory of polymer brushes. On the other hand, polymer stars form when linear polymer chains connected to one central monomer by one end. The chains stretch in radial direction because of the steric repulsion between monomers of nearby chains. Star-like structures reside in synovial fluid, blood flow and so on as macromolecular inclusions. It is known that macromolecular inclusions suppress mechanical instabilities in synovial joint^4^. In blood flow, there are macromolecular structures with architectures very similar to polymer stars. These star-like macromolecular structures form a gel-like network turning blood into clot. Here, I employ the density functional theory framework to extract equilibrium properties of a brush and a star interpenetrating into each other. My discussion continues by making a brief derivation of building blocks of the density functional theory framework. Then I describe equilibrium properties of brush and star, separately. Ultimately, I tackle the problem of interpenetrating brush and star through the DFT framework.

**Figure 1:**
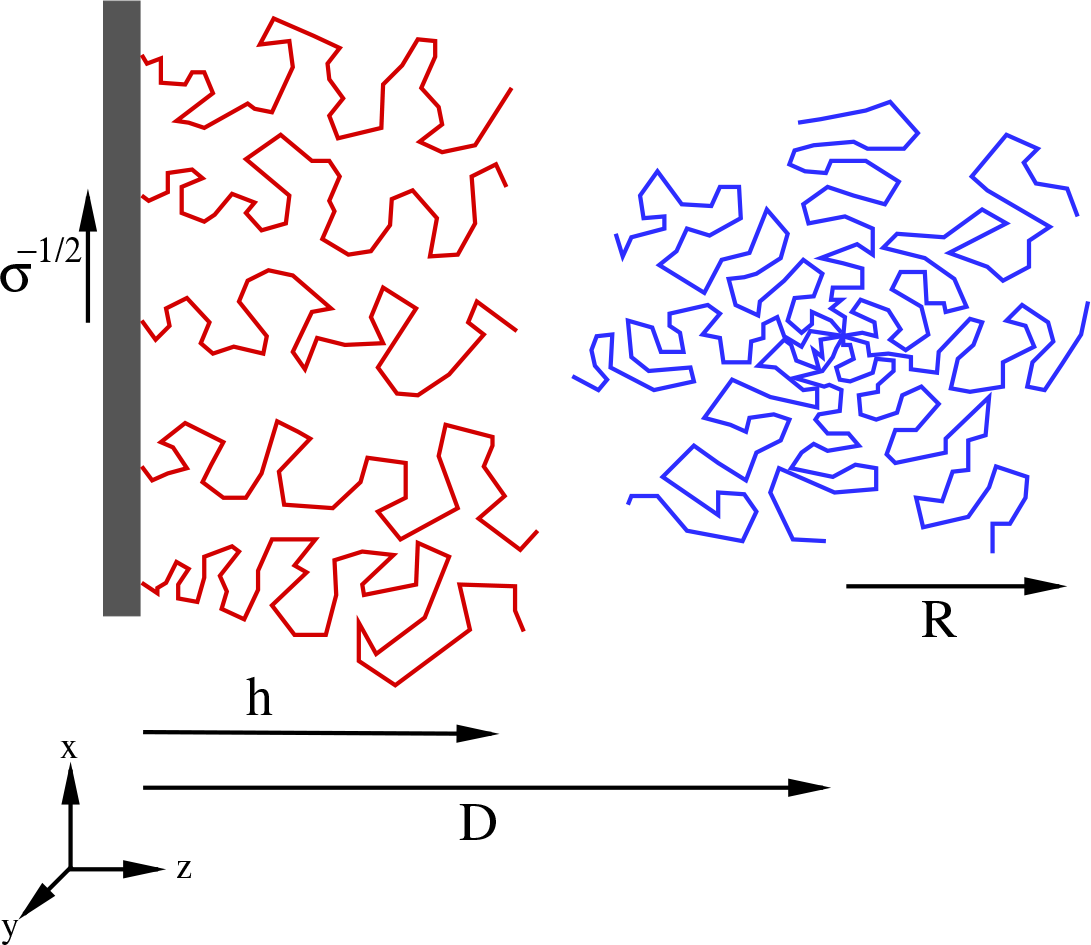
A two dimensional sketch of the problem. A brush with height *h* is grafted to a substrate at (*z* = 0) and a star with radius *R* is located at (*z* = *D*).

## DENSITY FUNCTIONAL THEORY (DFT) FRAMEWORK

### Polymer chain

A polymer chain with *N* monomers each of size *a* can be formulated through statistical mechanics as *N* freely rotating vectors in space. In order to obtain intrinsic properties of polymer chain, one needs to look at that chain under influence of an external force, say *f*. The Hamiltonian for such a system is given as follows,

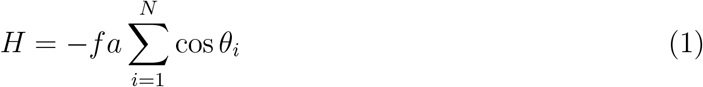

where *θ*_*i*_ represents the angle between any vector and the external force, which I take it to apply in the z direction. Then, the partition function is given as integration over all possible configurations of the vectors, which can only rotate leading to the following integral to be solved,

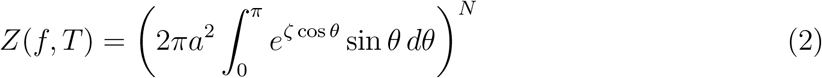

where *ζ* = *af/k*_B_*T* is a dimensionless quantity. The above integral can be exactly calculated giving to the following answer,

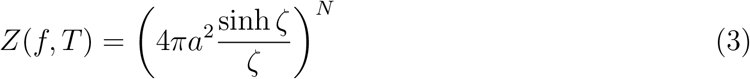

Then the next step would be calculation of the canonical free energy from the partition function leading the following answer,

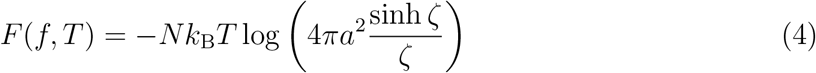

The first quantity which is interesting in polymer science is the expectation value of chain length. It turns out that in presence of an external force, the expectation value of the chain length does not vanish in the direction of the external force (i.e. z direction here) and it converges to the following answer,

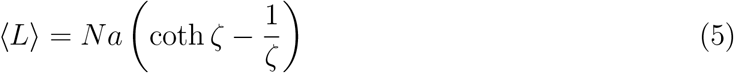

However, the mean square length is shown to be independent of the external force and has the following form,

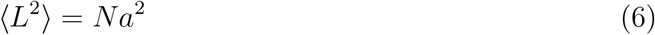

This is the prediction of the so called *Freely Jointed Chain* model which tells us that a polymer chain has a mean-square-length proportional to *N* ^1/2^. Another quantity which is interesting to think about would be the fluctuations in mean-square-length. One can calculate that as 〈*L*^2^〉 − 〈*L*〉^2^ leading to the following answer,

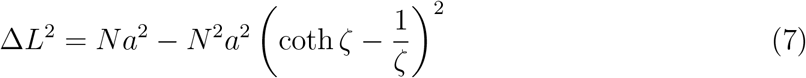

The above answer reveals a sophisticated behavior of Δ*L*^2^ in terms of parameters of the system which I discuss later. The elastic properties of the chain can be estimated by calculating the *susceptibility* which is defined as *κ* = −∂^2^*F* (*T, f*)/∂*f* ^2^. That would give us an answer of the following form,

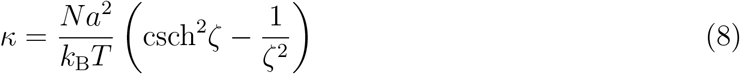

Since, the chain statistics at fixed external force ensemble requires that the chain length fluctuates, and on the other hand, I am interested in the statistics of a chain at fixed length, the *Helmholtz free energy* is my favorite one. In order to calculate that, one needs to make the transformation as *A*(*L, T*) = *F* (*f, T*) + *f* 〈*L*〉. Then, one gets the following answer for the Helmholtz free energy,

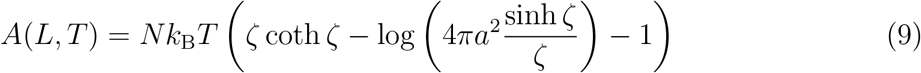

A good test of the chain statistics that have been obtained above would be to look at their asymptotic behaviors. For instance, the behavior of a chain when *ζ* approaches infinity. If the external force is much greater than the thermal energy one gets the following limiting answers,

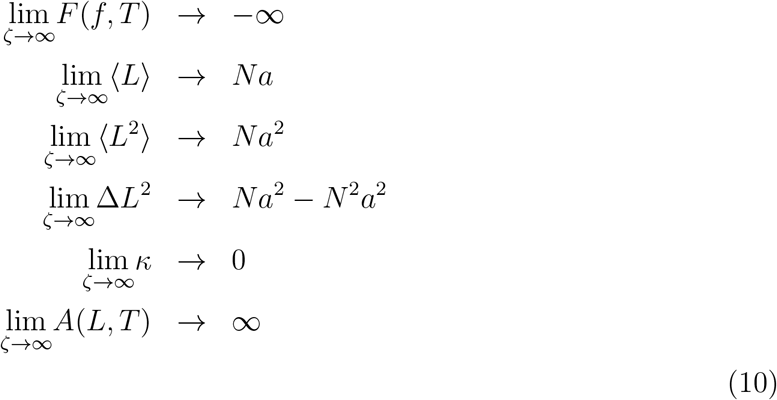

The above results tell us, correctly, that when a chain is subject to a large external force, it stretches to a the rod limit in the direction parallel to the force. The chain looses its elasticity and its free energy blows up to infinity. In such a limiting case, the fluctuations in chain length becomes imaginary which means that it is irrelevant to have a rod fluctuates.

On the other hand, one may look at the asymptotic behavior of the chain when *ζ* approaches zero. That means the chain is subject to the external forces which are much weaker than thermal energy. Note here that, in most of the biological systems, this conditions dominate. In order to calculate the physical quantities at *ζ* → 0, one must make Taylor expansion of every quantity around *ζ* = 0. Then, it gives us all quantities expanded in powers of *ζ*, and eventually, one can pick up a leading order term that dominates other terms for each quantity. In the following, there are expansions of all quantities in powers of *ζ*. For instance, the canonical free energy is expanded over even powers of *ζ* which means that the leading order term is a quadratic term.

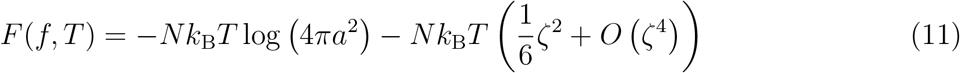

But the expectation value of the chain length is expanded over odd powers of *ζ*. That means there is a linear term as the leading order term here.

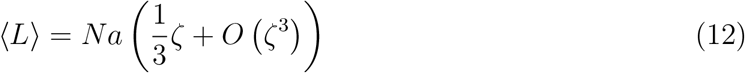

The power expansion of the susceptibility tells us a significant intrinsic property of the polymer chains. The susceptibility is expanded over even powers of *ζ*, however, there is a term independent of *ζ*. This term exists at *ζ* = 0 which means that it is an intrinsic property of the chain. Since, the compressibility is inverse of the susceptibility, one infers that compressibility of the chain equals 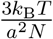
.

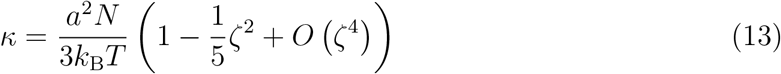

The power expansion of the chain length fluctuation indicates different scenarios that may happen.

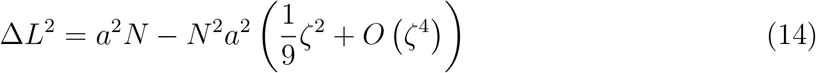

The chain fluctuation is expanded in even powers of *ζ* but there is a term independent of *ζ*. It tells us that in absence of the external force, the chain fluctuates dramatically, such that, the fluctuations equals *aN* ^1/2^ i.e. the whole chain region. Which is correct because monomers in a free chain actually can be located everywhere inside the coil. Nevertheless, as the external force increases, the fluctuations decrease until it vanishes at the rod limit.

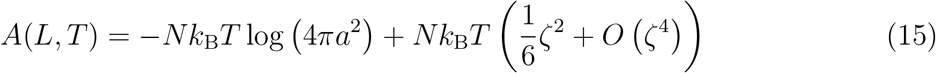

Now, if one looks at power expansion of the Helmholtz free, she/he would see that, in fact, the leading order term reads as *κ*^−1^〈*L*〉^2^/2. Actually, this term, which is quadratic in 〈*L*〉, describes all aspects of the chain at thermal equilibrium with a good accuracy.

### General short-range interacting monomers

After I introduced statistical properties of a polymer chain at thermal equilibrium, I would like to address a different system. Let us take *N* particles which can freely move in space and interact with each other through short-ranged forces. The Hamiltonian of such a system of *N* identical particles with short-ranged interactions is given as *H* = *K* + *U* with *K* total kinetic energy and *U* total potential energy of the system. The most simple case is system of *N* short-ranged interacting particles with two-body interactions. In such an particular case, total potential energy becomes a summation over all pair potentials *U*(***x***_*i*_, ***x***_*j*_) with ***x***_*i*_ and ***x***_*j*_ denoting position vectors of particle *i* and *j*. The total potential energy is a summation over *N* (*N* − 1)*/*2 possible pair potentials which excludes repeated and self interactions. The pair potential in systems that are exposed to no external force and with *translational invariance*, takes the form *U* (*r*) with *r* = |***x***_*j*_ − ***x***_*i*_|. In such an special case, one can write down the canonical partition function of the system as follows,

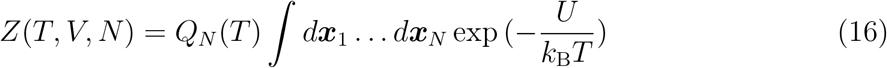

where *Q*_*N*_ (*T*) denotes the *kinetic integral* which contains integration over kinetic energy, and here, I do not take its details into account and I treat that as a prefactor. The integral in Eq. (16) is called *configuration integral* and contains information about interactions between particles. The partition function of interacting particles can not be solved exactly. However, when the system is dilute enough to take the particle number density as an small quantity i.e. *n* ≪ 1, one can make an evaluation to the partition function, free energy and the equation of state through expanding them in terms of *n*. To this end, we need to change the ensemble to the grand canonical ensemble where the particle number *N* is not fixed. The grand canonical partition function is given as follows,

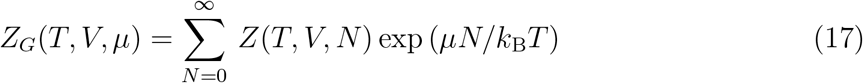

expanding Eq. (17) in powers of exp (*μN/k*_B_*T*) gives us the following series expansion,

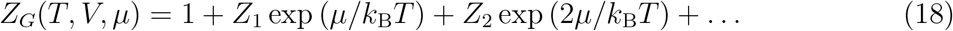

where for the sake of simplicity, *Z*_*N*_ denotes the canonical partition function with *N* particles. Having expanded the grand partition function in powers of exp (*μN/k*_B_*T*), using Φ(*T, V, μ*) = −*k*_B_*T* ln *Z*_*G*_(*T, V, μ*), one can calculate the grand potential as follows,

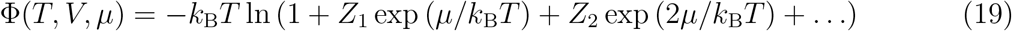

To expand the grand potential in powers of exp (*μN/k*_B_*T*), one can use a logarithm expansion as summation over the second terms to ∞ gives a quantity much smaller than unity. Therefore, we can make use of a logarithm expansion i.e. ln (1 + *y*) = *y* − *y*^2^/2 + *y*^3^/3 − … to expand the grand potential in powers of exp (*μN/k*_B_*T*) as follows,

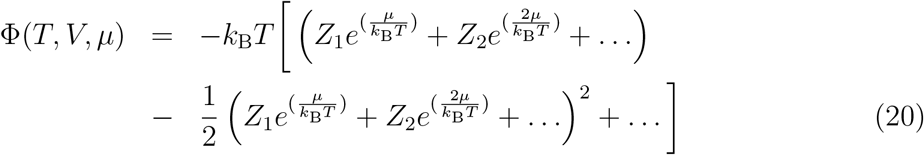

After simplifying Eq. (20), one may get the following power expansion for the grand potential,

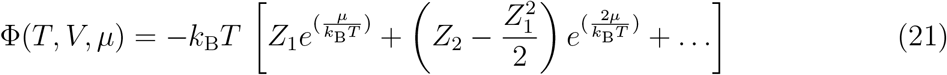

To expand the grand potential in powers of particle density, first of all, one needs to calculate the expectation value of particle number using *N* = −(∂Φ/∂*μ*)_*T,V*_ which is given as follows,

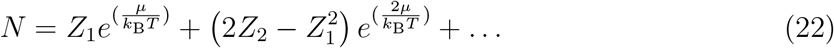

one must expand exp (*μN/k*_B_*T*) in powers of *N*, iteratively. To do so, one can insert Eq. (22) into an expansion of 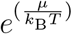 and determine unknown coefficients. This leads to an expansion which is given as follows,

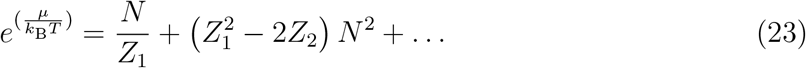

Inserting Eq. (23), into Eq. (21) and simplifying it one get the following expansion of grand potential per unit volume in powers of density,

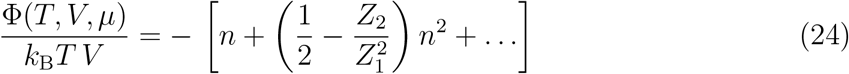

However, we need to know expansion of the Helmholtz free energy in powers of *N*. The Helmholtz free energy is obtained through Legendre transformation *A*(*T, V, N*) = Φ(*T, V, μ*)+ *μ N*. To find Helmholtz free energy in powers of *N*, one needs to expand grand potential as well as chemical potential according to Eqs. (23) and (24). At the end, Helmholtz free energy per unit volume is expanded in powers of *n* as follows,

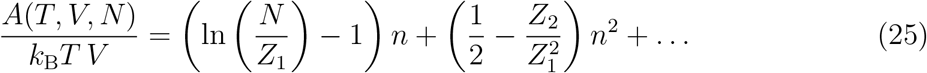

From Eq. (16), one finds that *Z*_1_ = *Q*_1_(*T*)*V* with *Q*_1_ = *a*^−3^ and *Z*_2_ is given as follows,

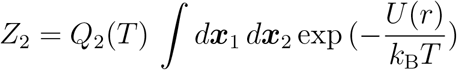

where *Q*_2_(*T*) = *a*^−6^/2. The configuration integral in the above form is not suitable as integrand contains only the pair potential. To make the configuration integral more suitable one needs to change coordinates to the centre of mass and relative displacement coordinates. This can be done simply by replacing *d****x***_1_ *d****x***_2_ with *d****x***_c_ *d****r*** with *d****x***_c_ a volume element in center of mass space. The integration over center of mass space gives us a total volume *V* as the integrand is independent of ***x***_c_. However, integration over the relative displacement depends upon pair potential.

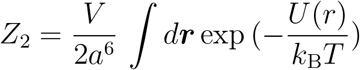

Eq. (25) is the *Virial* expansion of the free energy which one would write it down as,

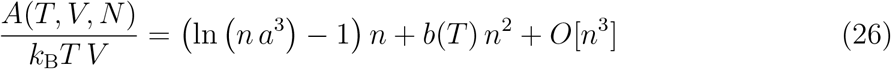

Coefficient of *n*^2^ in Eq. (26) is called the *second Virial coefficient* which is given as follows,

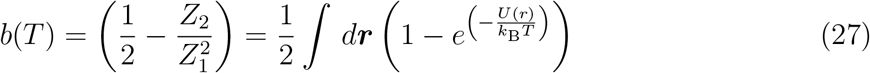

The second Virial coefficient is key quantity in describing short-ranged binary interactions between particles. For repulsive pair interactions (*U* (*r*) > 0), one finds *b* > 0, however, for attractive pair interactions (*U* (*r*) < 0), one finds *b* < 0. For hard spheres of diameter *a*, *b* = (2*π*/3)*a*^3^ and is independent of temperature. However, more realistic pair interaction which has been observed at molecular and atomic scales is the Lennard-Jones pair potential which reads as *U* (*r*) = 4*ϵ*[(*a/r*)^12^ − (*a/r*)^6^]. For a Lennard-Jones pair potential, one can calculate the second Virial coefficient at low temperatures as follows^3^,

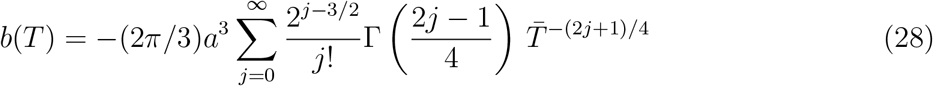

with *T̄* = *k*_B_*T*/*ϵ* defined as a dimensionless quantity^3^. For instance, if *T̄* = 1.68, then the above series converges to *b* = 4.19767.

In polymer science, a positive *b* refers to *good solvent* conditions and a negative *b* refers to *poor solvent* conditions. This definition is based on the fact that, monomers are treated much larger than solvent molecules and information about solvent molecules is contained in *b*, effectively. This type of solvent molecules treatment is called implicit solvent treatment. However, when solvent molecules are comparable with monomers, one has to take the excluded volume interactions between solvent and monomer through separate *b*. This type of solvent treatment is called explicit solvent treatment. Moreover, there is a certain temperature (*θ* temperature) in which *b* = 0. At this temperature, monomers refuse to make pair-wise interactions, however, the higher order interactions take place.

### Polymer brushes

Let us now employ the results of the previous sections to approach the problem of polymer brushes. One would build up the grand potential of the polymer brushes by adding the leading order term (which is quadratic in 〈*L*〉) of the Helmholtz free energy of a polymer chain Eq. (15) and the second term in the Virial expansion of the Helmholtz free energy of a short range interacting gas of monomers (which is quadratic in *n*) Eq. (26). But these are the Helmholtz free energy. In order to have the grand potential, one would make a transformation as Φ(*L, T, μ*)/*V* = *A*(*L, T, N*)/*V* − *μn* where *μ* is a Lagrange multiplier for the constraint of fixed number of particles (which is called the chemical potential).

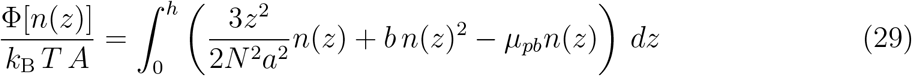

The above equation is the grand potential of the polymer brushes per *k*_B_*T* per unit surface area that has been proposed by Hirz^6,7^. Now, one would like to get to know what the density, brush height and the chemical potential are at thermal equilibrium. Those could be calculated through solving the following coupled equations.

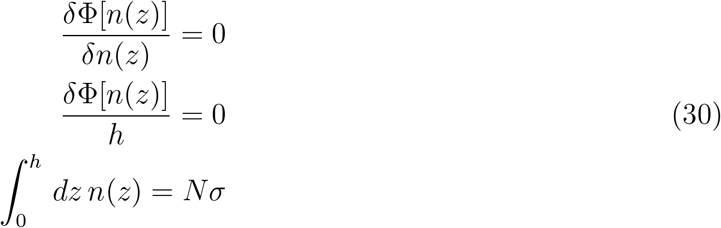

where *σ* would be the grafting density of the brush. The solutions of the above set of coupled equations are listed as follows,

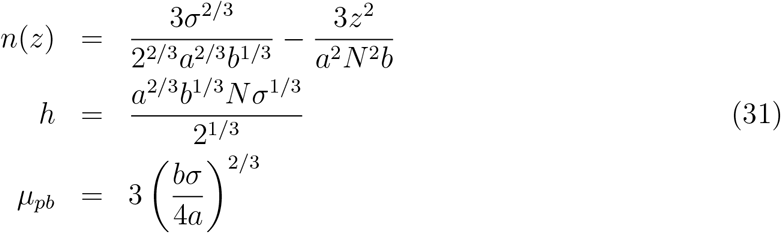

It turns out that the density profile is a parabolic function of z, and the brush height and the chemical potential are scaled by system parameters according to universal power laws.

Remarkably, the brush height scales as *N* which means that the chains strongly stretched in perpendicular direction. However, the chemical potential is independent of *N*. The total grand potential follows the universal power laws as follows,

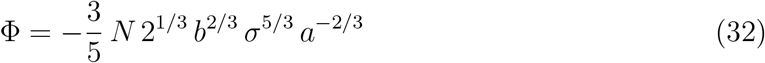

### Polymer stars

The polymer star is an spherically symmetric structure in which the distance from the center, i.e. radius *R*, determines the differences. Let us say, every where on a same radius have the same property. So, that tells us that the spherical polar coordinates is the most convenient set of coordinates one can work with. The grand potential functional for a polymer star with degree of polymerization *M* and functionality *f* (number of chains) could be written down as follows,

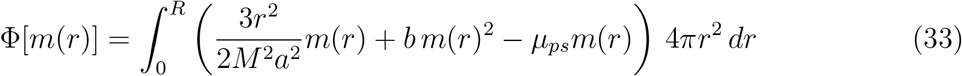

where *m*(*r*) is the density profile in radial direction and *μ*_*ps*_ the chemical potential. In order to find the density profile, radius and the chemical potential at equilibrium, one would need to solve the following set of coupled equations,

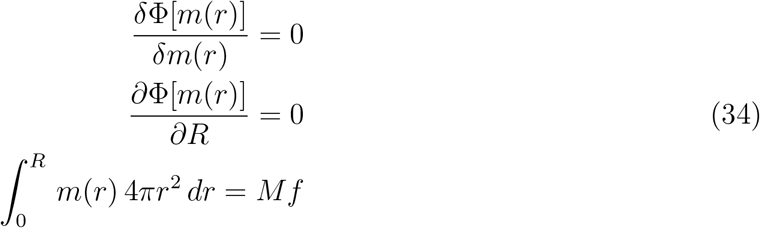

Those coupled equations are solvable exactly and the solutions are given as follows,

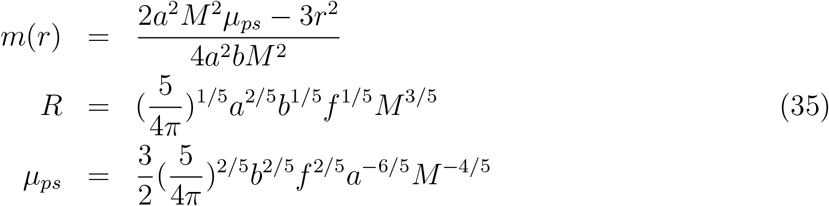

the density profile is parabolic in radial direction (similar to brush), the radius scales with degree of polymerization as *M* ^3/5^ (similar to a single chain) and it scales with functionality as *f* ^1/5^. Those solutions reveal the following universal power laws for the grand potential,

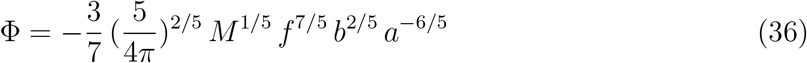

which strongly depends upon functionality *f* ^7/5^ and weakly upon degree of polymerization *M* ^1/5^. This tells us that the energy content of a polymer star is extremely sensitive (less than a brush) to number of chains in that. One would relate that to the spherical symmetry of the star and the fact that the chains are connected in a point. Doubling number of chains in an star would increase its energy by a factor of 2^7/5^, however, energy of a brush is increased by a factor of 2^5/3^. This may tell us that one can add more chains to a brush-like structure rather than a star-like structure and this is incredible fact.

## RESULTS

### Interpenetrating brush and star

Now, let us take a brush and an star a distance *D* away from each other. One would write down the grand potential for such a system as follows,

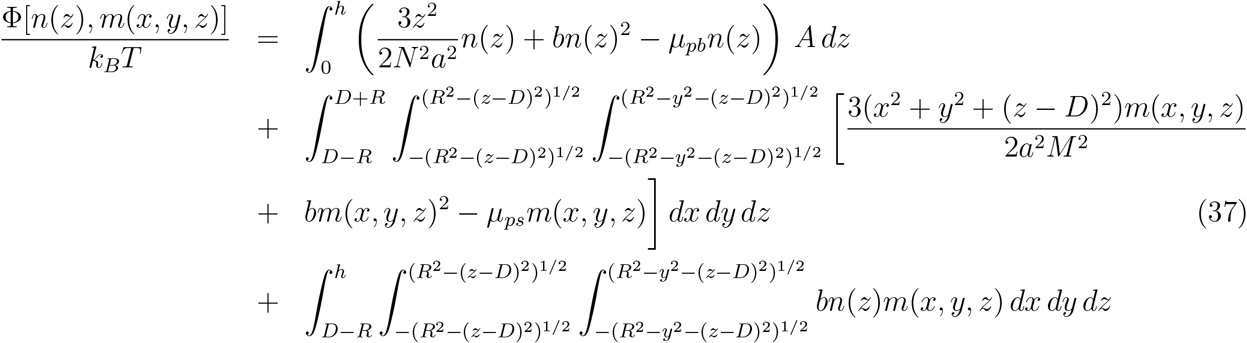

There is a point here that makes the spherical coordinates inappropriate. When star enters the brush, the interpenetration region has a shape of a cup (as long as the asphericity of the star maintained at small interpenetrations). That is why one needs to translate the grand potential of the star in the Cartesian coordinates. To do so, the integration limits are determined by *R*^2^ = *x*^2^ + *y*^2^ + *z*^2^.

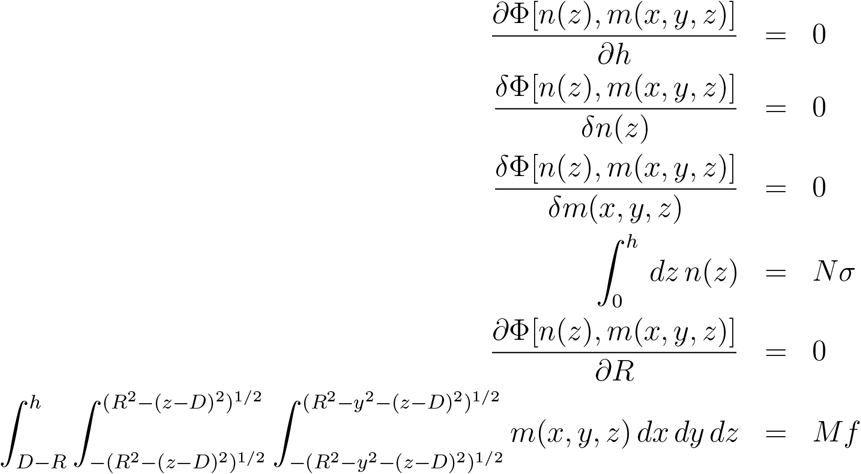

The above set of coupled equations have to be solved in order to have density profiles, brush height, star radius and chemical potentials. The equilibrium density profiles are given as follows,

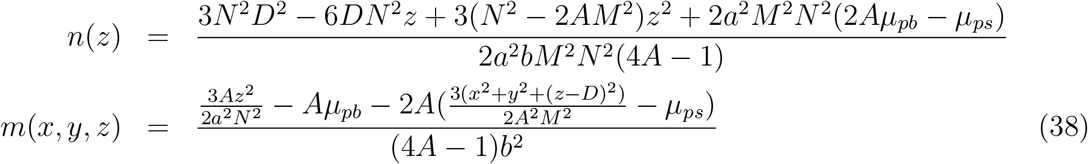

A sample plot of the above density profiles are plotted in Fig. (2) by choosing the system parameters as *N* = 30, *a* = 1, *b* = 2.09, *σ* = 0.1, *M* = 15, *f* = 10 and *D* = 19. It turns out that the density profile of the brush is not influenced that much by star. However, star swells and repelled by the brush. Two effects that are expected already. Fig. (2) reveals that for a given *D* the star’s position for interpenetrating and non-interpenetrating systems is different. It turns out that interpenetrating star stays two or three segment lengths further away from the brush. Obviously, this difference is created as a result of large density fluctuations within the interpenetration region.

**Figure 2:**
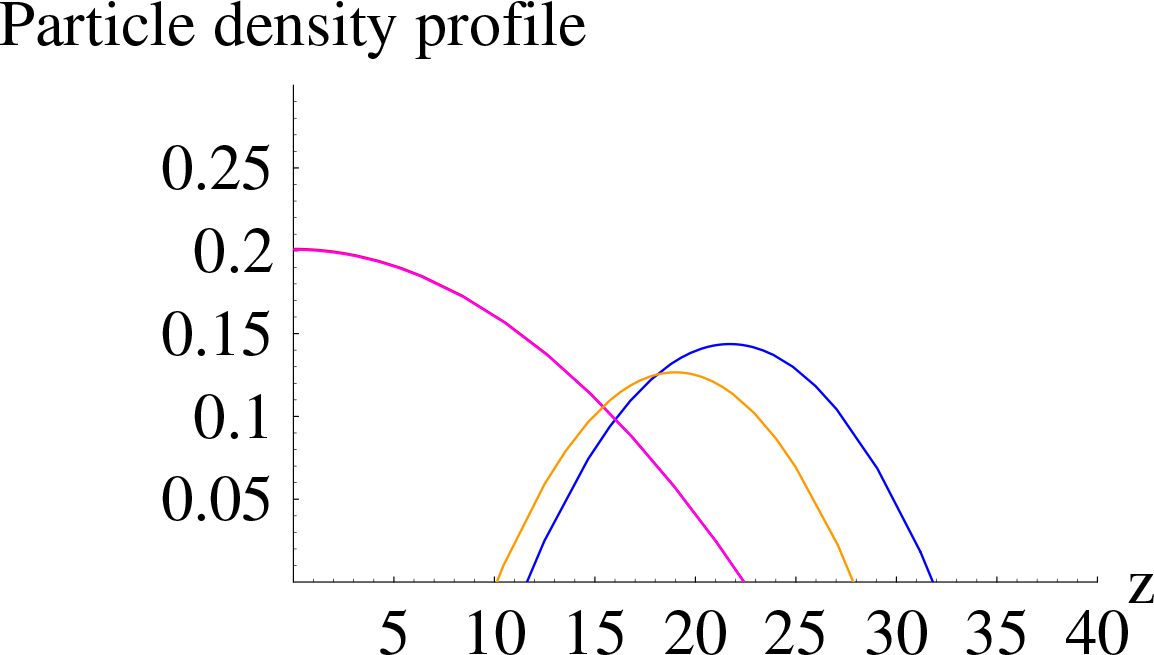
Density profiles of the brush and the star in the direction perpendicular to the brush.

The set of coupled equations in Eq. (38), is neither analytically nor numerically solvable. In order to calculate, the brush height, the star radius and the chemical potentials, one needs to employ the *RootFinding* package of the *Mathematica* software. Fig. (3), shows the brush height *h* and the star radius *R* as functions of *D*. These plots tell us that when star sucks into the brush, its radius decreases, though, the brush height does not change.

**Figure 3:**
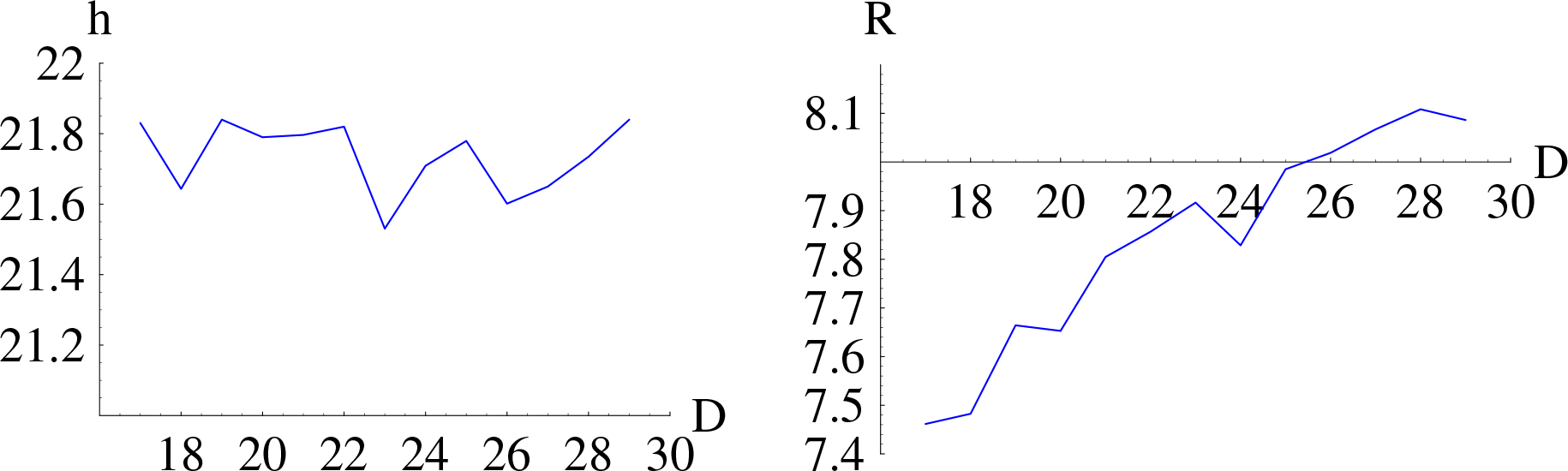
The brush height (left) and the star radius (right) as functions of *D*.

## CONCLUDING REMARKS AND DISCUSSION

The interpenetration between different types of macromolecular structures demands an extremely non-trivial theoretical effort. As Edwards’s equation has been proposed to take the steric repulsion between polymers into account by introducing a self-consistent-field theory approach, however, that does not work even for the simplest cases. The reason would be the increasing complexity associated with calculating the integrals encountered. What I would like to show here was that the DFT framework works for almost any polymeric structures.

I showed that the star is repelled by the brush as it interpenetrate with brush. At the same time, the star collapses by further sucking into the brush. Well, it has been observed in MD simulations that an star undergoes a discontinuous absorption transition by the brush when they are oppositely charged^9^. The existence of such a spatial difference between interpenetrating and non-interpenetrating star position, tells us that if an star is going to suck into the brush, then it supposed to overcome a barrier. That is why a discontinuous absorption transition has been observed there^9^. Another remarkable prediction of the current study would be the collapse of the star by further sucking into the brush. That could happen when the star, which is spherically symmetric, interpenetrate with the brush, which has parallel chains. Then the star collapses to minimize large steric repulsion between its arms and the brush chains. The star can maintain its asphericity only at very small interpenetrations. Larger interpenetrations require the star to loose its asphericity based on the energy minimization principle.

The most useful aspect of this study would be to take the simultaneous deformations of the brush and star into account by solving a set of coupled equations. This allowed approaching one of the most non-trivial problems in theoretical polymer science. The same DFT framework could be employed to tackle more sophisticated problems such as interpenetrating stars or denderimers and even the long range interactions can be taken into account later. For instance, interpenetrating Polyelectrolytes with different architectures could be interesting.

## References

1. Rubinstein, M.; Colby, R. H. Polymer Physics; OUP Oxford, 2003.

2. Advincula, R. C.; Brittain, W. J.; Caster, K. C.; Rühe, J., Polymer Brushes; Wiley-VCH, Weinheim, 2004.

3. Schwabl, F.; Brewer, W. D. Statistical Mechanics, Advanced Texts in Physics; Springer Berlin Heidelberg, 2006, 313–316.

4. Kreer, T., Polymer-brush lubrication: a review of recent theoretical advances; Soft Matter 2016, 12, 3479.

5. Gennes, P. G., Scaling Concepts in Polymer Physics; Cornell University Press, 1979, 74–75.

6. Hirz, S. J., Modeling of Interactions Between Adsorbed Block Copolymers; Master Thesis, University of Minnesota, Minneapolis, MN, 1988.

7. Safran, S., Statistical thermodynamics of surfaces, interaces and membranes, Boca Raton, Israel, 2003.

8. Farzin, M., Polymer brush bilayers at thermal equilibrium: A theoretical study, 2018, https://doi.org/10.1101/316141.

9. Farzin, M.; Kreer, T.; Sommer, J.-U., Absorption of a Polyelectrolyte star by an oppositely charged Polyelectrolyte brush, Unpublished.

